# Top-Down Ion Mobility Separations of Isomeric Proteoforms

**DOI:** 10.1101/2022.07.23.501273

**Authors:** Francis Berthias, Hayden A. Thurman, Gayani Wijegunawardena, Haifan Wu, Alexandre A. Shvartsburg, Ole N. Jensen

## Abstract

Continuing advances in proteomics highlight the ubiquity and biological importance of proteoforms - the proteins with varied sequence, splicing, or distribution of post-translational modifications (PTMs). The preeminent example is histones, where the PTM pattern encodes the combinatorial language controlling the DNA transcription central to life. While the proteoforms with distinct PTM compositions are distinguishable by mass, the isomers with permuted PTMs (“localization variants”) commonly coexisting in cells generally require separation before mass-spectrometric (MS) analyses. That was accomplished on the bottom-up and middle-down levels using chromatography or ion mobility spectrometry (IMS), but proteolytic digestion obliterates the crucial PTM connectivity information. Here we demonstrate baseline IMS resolution of intact isomeric proteoforms, specifically the acetylated H4 histones (11.3 kDa). The variants with a single acetyl moiety on five alternative lysine residues (K5, K8, K12, K16, K20) known for distinct functionalities in vivo were constructed by two-step native chemical ligation and separated using trapped IMS at the resolving power up to 350 on the Bruker TIMS/ToF platform. Full resolution for several pairs was confirmed using binary mixtures and by unique fragments in tandem MS employing collision-induced dissociation. This novel capability for top-down proteoform characterization is poised to open major new avenues in proteomics and epigenetics.

---

The fully sequenced human genome features about 20,000 genes, yet human bodies comprise millions of protein species. That is, each gene engenders a family of proteoforms with alternative splicing, combinations of post-translational modifications (PTMs), or minor amino acid sequence deviations.^1^ Protein diversity varies widely across the cell types in individuals and over the population, and responds flexibly to various external events and physiological/disease states.^2^ Mapping, understanding, and eventually predicting these networks and processes has emerged as the defining frontline of biology.^3^ Perhaps most complex and also consequential case is histone proteins found in all eukaryotic cells.^4^

All molecular events therein involving DNA (replication, transcription, repair, and recombination) are related to dynamic changes in the chromatin in response to events such as exogenous stimuli or cell-specific developmental programs.^5^ The DNA compaction, accessibility and genome regulation are regulated in nuclei by several histone proteins (~100 - 130 residues), specifically the combinatorial pattern of their PTMs such as methylation (me), trimethylation (me3), acetylation (ac), and/or phosphorylation (p)^6^ - the “histone code” controlling the higher-order chromatin structure.^7^ Measuring the full complement of expressed proteoforms and linking each to the ensuing 3-D structure and biological outcomes is a most staggering analytical and biomedical challenge.^8^ Of the >10^4^ potential histone forms, only ~1000 have been discovered so far.^9^ Even in that minimal set, many are structural isomers (variants) with differing PTM localization. While PTMs cluster on the N-terminal tails (~50 residues) that protrude from the nucleosome, some with important functions reside in the core domain. Hence, whole intact histones must be characterized.

Proteoforms are normally identified by unique fragments observed in tandem mass spectrometry. While non-ergodic methods such as electron capture or transfer dissociation (ECD/ETD) that sever the protein backbone but not PTM links are preferred, standard collision-induced dissociation (CID) is suitable with tightly bound PTMs such as ac.^10–12^ However, only the two variants with distal PTMs yield unique products, obstructing the characterization of real samples including many more actual or potential variants. Hence, effective separations prior to MS analyses are required. Liquid chromatography (LC) could resolve some variants for small tryptic peptides (~10 residues) from histones in bottom-up workflows, but not large peptides in middle-down analyses or intact proteins.^13,14^ The problem is that any enzymatic digestion deletes the crucial information on PTM connectivity and cross-talk.^15^

A growing alternative to LC is post-ionization separation in gases using ion mobility spectrometry (IMS), comprising linear IMS based on the absolute ion mobility (*K*) at normally moderate electric field intensity (*E*) and differential or field asymmetric waveform IMS (FAIMS) that captures the increment of *K* at high *E* levels.^16–18^ Both methods are now implemented in commercial IMS/MS platforms and increasingly adopted in bottom-up proteomics. Both are based on the 3-D ion geometry and, for FAIMS, its reversible evolution (generally expansion) during the heating in high-field periods. As the PTM localization affects the macromolecular folding and its thermal stability, variant peptides or proteins ought to be separable. Indeed, both linear IMS and FAIMS have broadly separated variant histone N-tail domains and, in a major advance over LC, complete N-tails with me, me3, ac, or p on alternative sites. The separations of intact proteins within the top-down paradigm are still needed to comprehensively map the PTM patterns, confidently identify and quantify variants across the dynamic range, and unambiguously elucidate their distinct biological functions.

The IMS analyses of isomeric histone tails and other peptides of similar size (~5 kDa) have shown the pressing demand for resolving power, partly as the conformational multiplicity for each variant impedes their separation.^19–25^ The expanding number and diversity of coexisting energetically competitive conformations for longer peptides suggests the need for yet greater resolution with proteins. Over the last decade, the novel dynamic-field IMS methods including trapped IMS (TIMS),^26–28^ structures for lossless ion manipulations^29^ and related cyclic IMS,^30^ and high-definition FAIMS^31^ have raised the IMS resolving power (*R*_IMS_) by order of magnitude, up to ~300 routinely and ~1000 in selected demonstrations.^32–34^ This remarkable advance now embodied in commercial instruments enables taking the separations of isomeric proteoforms to the intact-protein realm.

Here, we explore the separations of whole histone variant isomers generated by nanoelectrospray ionization (nESI) source using TIMS.^26–28,35^ The pull of ions radially confined by rf in an electrode tunnel into the time-of-flight (ToF) mass spectrometer by gas flow is opposed by retarding dc field gradient.^28,35^ Species with different *K* values are balanced at different points along the tunnel and sequentially eluted by ramping the field down. We look at five biologically pertinent H4 species monoacetylated on K5, K8, K12, K16, or K20. These modifications are implicated in transcriptional regulation, with the gene repression for K20 but expression for others^36,37^ showcasing the importance of exact PTM position. Substantial resolution of multiple variants across the charge states, validated by CID upon mobility selection, demonstrates the broad capability for isomeric proteoform separations within top-down workflows.

## EXPERIMENTAL METHODS

### Synthesis of histones and sample preparation

The acetylated H4 variants (102 residues) were assembled via native chemical ligation^38^ from three pieces built by the regular Fmoc protocol: (I) N-terminal segment H4(1-37)-NHNH_2_ with ac on the defined lysine (K5, K8, K12, K16 or K20), (II) middle segment Tfa-Thz-H4(39-75)-NHNH_2_, and (III) C-terminal segment Cys-H4(77-102)-OH (Supplemental Figure S1). The trifluoroacetyl (Tfa) group in II protected the thiazolidine (Thz) from oxidation during the hydrazide activation step.^39^ Upon the hydrazide-based ligation^40^ of (II) and (III), the Tfa deprotection and Thz-to-Cys conversion were successively performed in “one-pot”. That (II + III) construct was joined to a (I) variant to yield a cysteine-containing H4 sequence, which was desulfurized to produce the final protein (details in SI). Those were purified by preparative HPLC and validated by accurate mass (average *m* = 11,278 Da) using a nESI/Orbitrap Elite platform (Thermo). The individual variants and binary mixtures were dissolved to 5 μM in the 50:49:1 methanol/water/formic acid solvent. This extremely acidic (pH = 2.2) organic medium shifts the ESI output to unfolded less conformationally heterogeneous peptides at high charge states (z) for improved IMS separation.^19–25^

### IMS/MS/MS analyses

We employed a timsTOF Pro2 hybrid Q-TOF tandem mass spectrometer (Bruker Daltonics). The principle of TIMS separation, explained above, is illustrated in Figure 1. Samples were infused to the Captive Spray nESI emitter at a flow rate of 0.3 μL/min using a nanoElute LC system. The *m/z* scale was calibrated using the ESI-L low concentration tune mix (Agilent). As the dynamic-field IMS systems including TIMS cannot measure the absolute *K* values *a priori*, a calibration using species with established mobility is required. For that, we deposited on the ESI source air filter two species from the above tune mix -at *m/z* of 622.03 (1/*K*_0_ = 0.992 Vs/cm^2^) and 922.01 (1/*K*_0_ = 1.12 Vs/cm^2^) where *K*_0_ is the reduced mobility. The calibration preceded each analysis and followed all TIMS setting changes. The ion-neutral collision cross sections with N_2_ gas (CCS) are derived from mobilities via the Mason-Schamp equation and calibrant CCS values (203.0 and 243.6 Å^2^, respectively).

**Figure 1.**
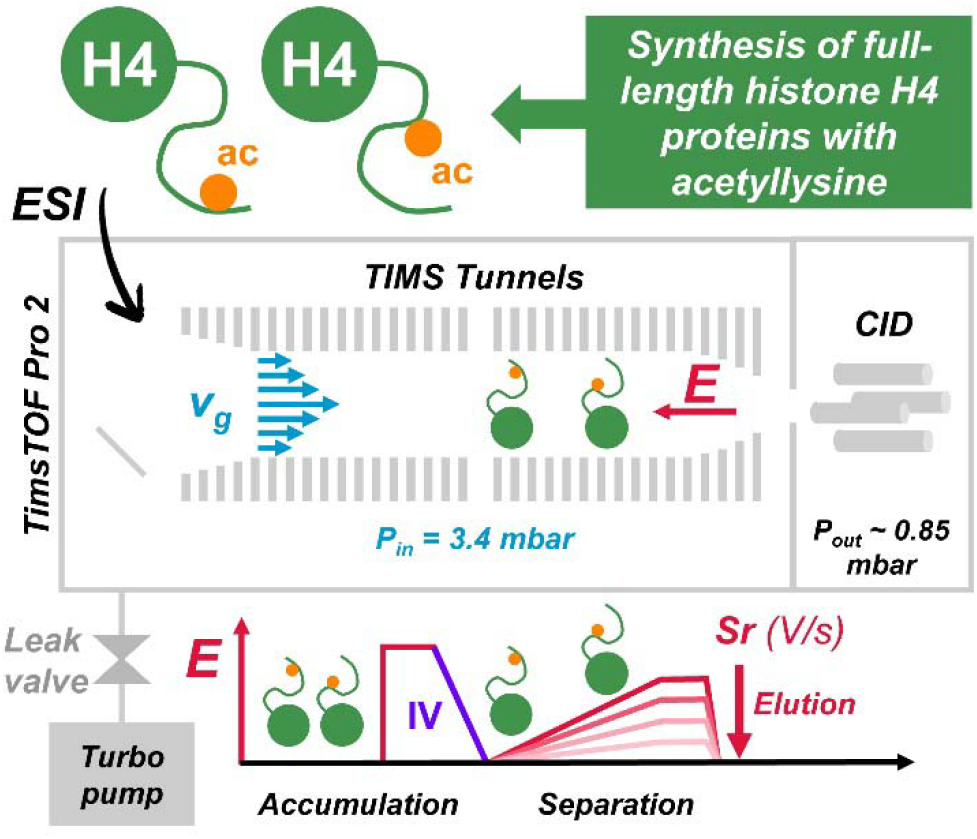
Schematic representation of TIMS separation of PTM positional isomers of intact histone H4.

Again, separating large macromolecules with minute structural distinctions necessitates the ultimate resolving power. That for TIMS (defined as *K*_0_/*w*, where *w* is the fwhm peak width in mobility spectrum) equals:

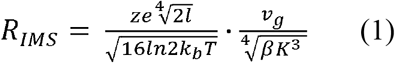

where *v*_g_ is the gas flow velocity governed by the difference between pressures at TIMS stage entrance and exit (*P*_in_ and *P*_out_), β is the field ramp rate, and z, e, l, k_b_, T, K are the charge of the ion, the elemental charge, the length of the electric field gradient plateau, the Boltzman’s constant and the ion mobility coefficient, respectively. We have found the maximum *R*_IMS_ at *P*_in_ = 3.4 mbar and *P_out_* = 0.85 mbar, by regulating the vacuum through a leak valve located at the entrance stage. As the β value is inversely proportional to the ramp time *t* and increases for wider scanned *K*_0_ ranges, we pushed up *t* and minimized those ranges by focusing on narrow windows containing the species of interest. Upon trying *t* up to the instrumental maximum of 2.0 s in the present experimental conditions, we achieved the best overall performance at *t* = 1.3 s with the voltage ramp rate of *Sr* = 17 V/s. At longer times, the signal dropped markedly with no significant or consistent resolution gain (perhaps because of poor ion count statistics): not surprising as *R*_IMS_ scales only as β^−1/4^ (eq 1). The initial wide-range acquisitions to pin the windows for targeted scans were faster (*Sr* = 86 V/s).

A crucial parameter in IMS analyses of fragile or flexible species is the injection voltage (IV) into IMS cell that controls the isomerization driven by transient collisional heating. We have found the instrumental maximum of 200 V to narrow the peaks (in line with prior work) and generally maximize resolution as the diverse kinetically trapped geometries anneal to common low-energy basins. Energetic injection into IMS cells (including TIMS) can also fragment ions,^41^ but that normally applies only to smaller precursors and was not encountered here. The species resolved by TIMS were selected for CID in N_2_ gas with the *m/z* isolation window of 5 units and excitation voltage of 30 V.

## RESULTS AND DISCUSSION

### TIMS separation of intact histone proteoform isomers

nESI mass spectra were similar for all isomers of protonated H4ac with sufficient signal at *z* = 10 - 19 (Figure 2a). As is usual, the ion mobilities increase at higher z sublinearly because protein unfolding elevates the CCS values (Figure 2b). Most H4ac variants exhibited multiple conformers for each charge state, which often leads to overlapping mobility peaks. Nonetheless, the spectral differences between some H4ac isomers allowed for their separation. Those come out in narrow range scans as reduction of *Sr* by 5× properly augments *R*_IMS_ by ~50% (Figure 2c - f). We pursued more detailed TIMS analysis of binary mixtures of H4ac isomers at the optimum charge states *z* = 16 - 19. Average R_IMS_ value increases with the number of charges from ~185, ~225, ~250 to ~260 for z = 16, 17, 18 and 19, respectively. We define full resolution as the ability to separate a H4ac isomer at >50% of its maximum intensity from another isomer and with <10% interference from an overlapping isomer). We define partial resolution as observing at least one isomer with <20% overlap from other isomers.

**Figure 2.**
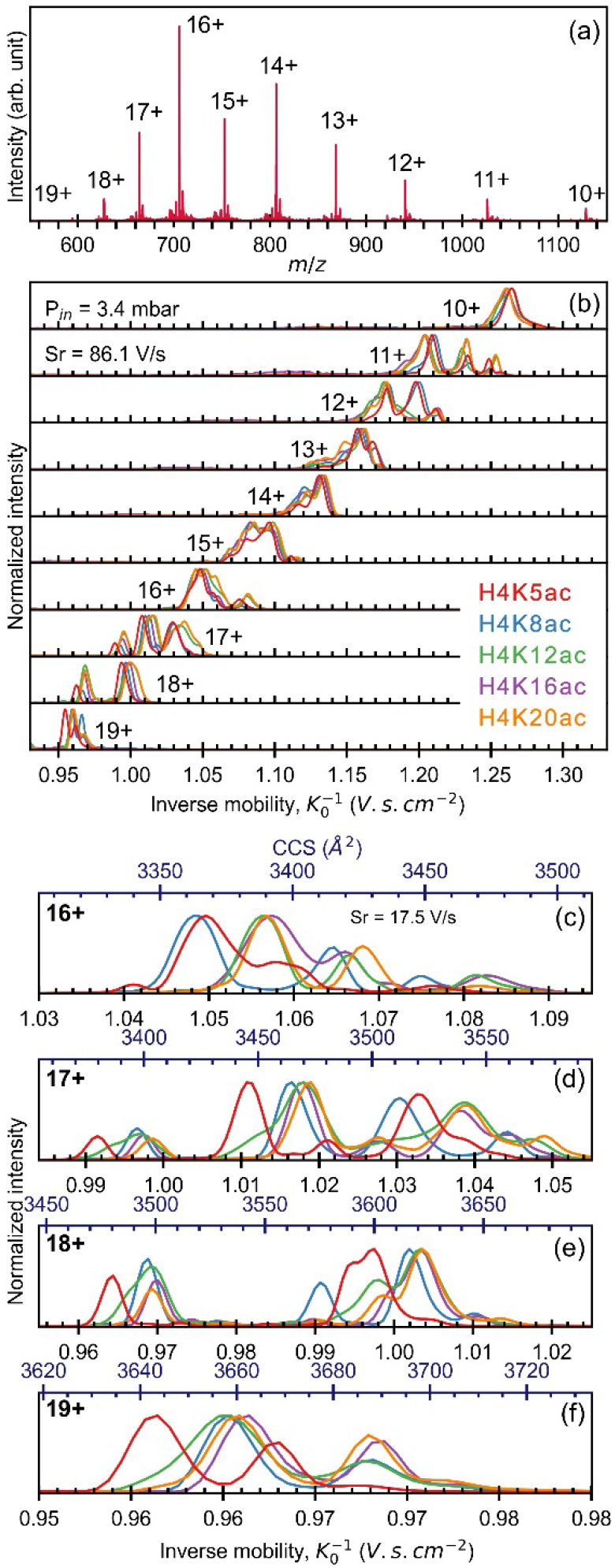
Performance of TIMS separation for each individual positional isomer of H4ac; (a) mass spectrum of protonated intact histone H4K5ac; (b) broad survey of TIMS separation for H4K5ac (red), H4K8ac (blue), H4K12ac (green), H4K16ac (violet) and H4K20ac (orange) with Sr = 86.1 V/s for z = 10-19+; (c), (d), (e) and (f) high resolution TIMS spectra as a function of K_0_^-1^ (bottom axis) and CCS values (top axis) for the charged states z = 16-19+.

Full TIMS separation is achieved for the isomeric intact H4ac pairs K5/K8, K5/K16, K5/K20, and K8/K12 (Figure 3). Partial resolution is achieved for K5ac/K12ac, K8ac/K16ac and K8ac/K20ac, respectively. Despite minor interferences and for z = 18, K8ac can be filtered from K16ac and K20ac for K_0_ ~ 0.88 Vs/cm^2^, but the converse is not true. Additional TIMS spectra for isomeric pairs of H4ac proteoforms and charge states are provided in Supplemental Figures S2 and S3. Of all the proteoforms analyzed, K5ac is the one that stands out the most from the others, followed by K8ac and K12ac. Separation, highlighted by colored dashed lines on Figure3, occurs on several distinct mobility values and for different charge states, thus multiplying the probability of filtering a particular variant in the case of more complex mixtures. Resolving power, average inverse mobility and collision cross section values of each variant are given in Supplemental Table S1 for z = 16 – 19. Among the 10 different pairwise combinations, three are fully separated and four are partially separated. K16 and K20 remain indiscernible to each other, and to K12, at the precursor ion level.

**Figure 3.**
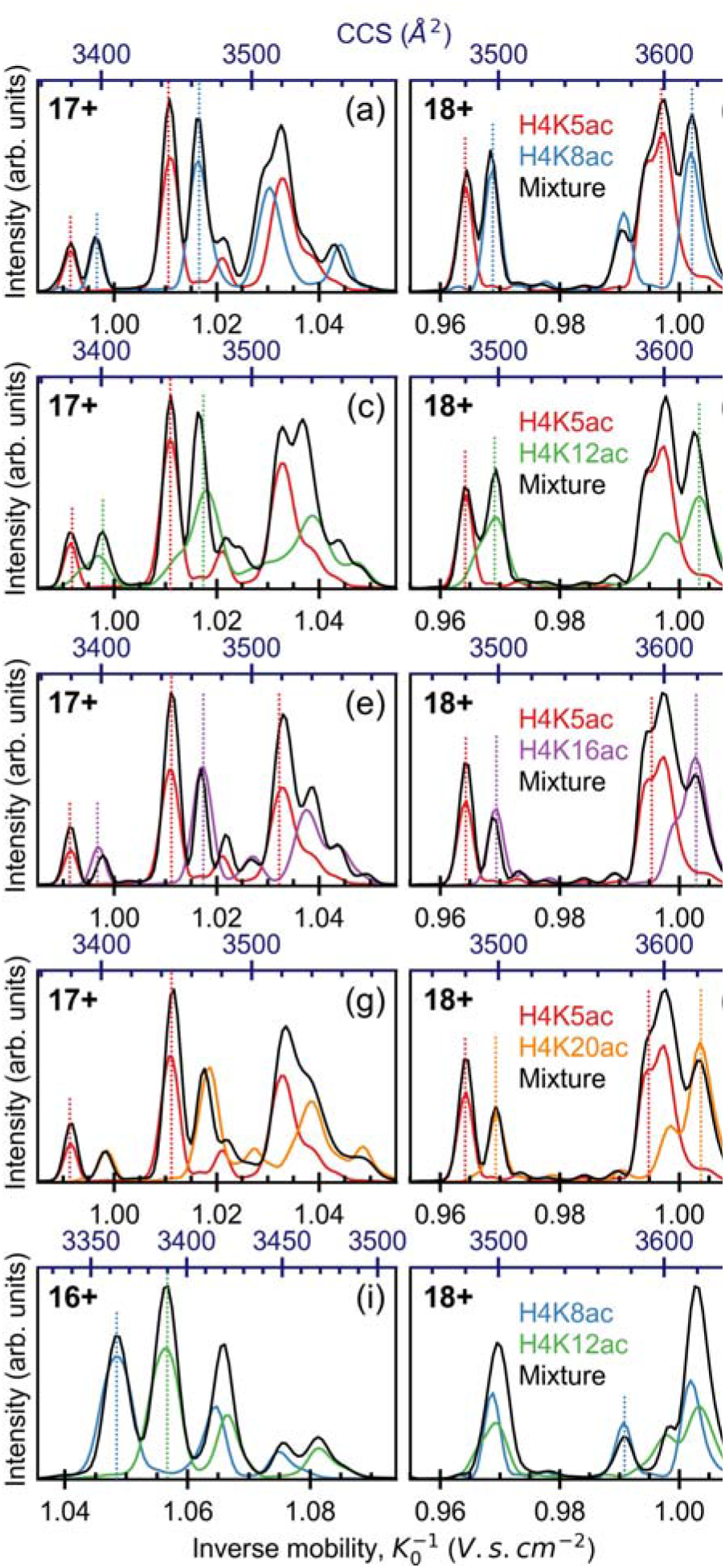
TIMS-MS separation of binary mixture (black) of H4K5ac (red), H4K8ac (blue), H4K12ac (green), H4K16ac (violet) and H4K20ac (orange) for different charge state. Experiments were performed using P_in_ = 3.4 mbar and an electric field scan rate of 17 5 V/s

### Isomeric histone proteoform assignment at MS/MS level

Product ions produced by CID MS/MS assesses the identity of each proteoform in the mixtures. Fragments of type b were favored over the y-fragments due to the absence of isobaric overlap by means of their lighter mass and lower charge number. Informative ions b_5_^2+^, b_8_^2+^, b_12_^3+^ and b_17_^4+^ are detected in sufficient amount to confidently identify each proteoforms of H4ac (Figure 4a at z = 18). By way of example, selected TIMS MS/MS spectra of binary mixtures are presented in Figure 4b-d. As the CID event occurs after the TIMS separation, the informative MS/MS fragment (color lines in Figure 4b-d) of each variant exhibits a similar mobility distribution to that of its precursor. Unique fragments do not only validate the TIMS separation without the use of reference compounds (Figure 4b and c), but it expands the performance of the workflow by disentangling unresolved variants at TIMS-MS level as illustrate for the K16/K20 pair (Figure 4d).

**Figure 4.**
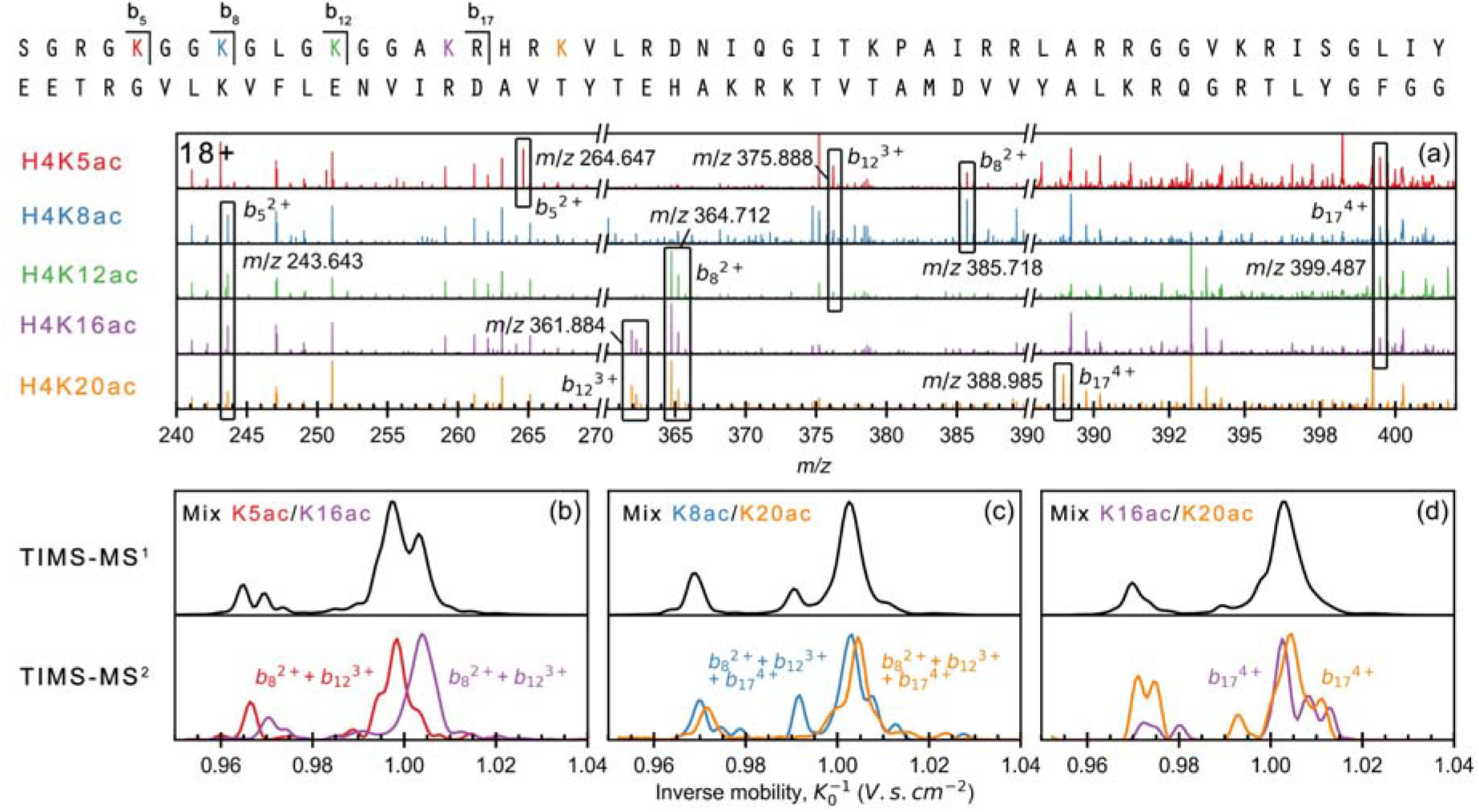
Separation and identification of acetylated H4 proteoform isomers by TIMS-CID MS/MS. (a) CID MS/MS fragment ion spectra generated from intact acetylated H4 proteoform isomers at charge state 18+; TIMS-CID MS/MS spectra of binary mixture H4K5ac/H4K16ac (b) H4K8ac/H4K12ac (c) and H4K12ac/H4K20ac (d). In (b)-(c) panels, the TIMS-MS spectra of the precursor ion (z = 18) are plotted in black. The informative fragments of the different positional PTM isomers of H4 are plotted in red (H4K5ac), blue (H4K8ac), violet (H4K16ac) and orange (H4K20ac) lines.

## CONCLUSIONS AND OUTLOOK

We report the first separation of isomeric intact proteoforms. The topical monoacetylated H4 histone variants (constructed by native chemical ligation) were resolved by ion mobility and identified by tandem MS within a top-down workflow implemented on the Bruker TIMS/TOF platform. Such separations require the ultimate IMS resolving power (*R*_IMS_) to disentangle subtle folding differences arising from PTM transposition within rich conformational ensembles prevalent for gas-phase protein ions. With extremely slow mobility scans to maximize resolution, the *R*_IMS_ for unfolded species in higher charge states *z* was up to ~350 and ~250 on average.

These metrics match those for small peptides and approach the instrumental limit of ~350,^20^ which means little peak broadening due to merged conformers. The peaks for lower z are broader, as diverse folded and unfolded geometries form the families of similar unresolved conformers. Such isolation of essentially single conformers is not seen in drift-tube (DT) IMS at similar instrumental R_IMS_, but mirrors that attained for high z in high-definition FAIMS.^42^ This shows the role of conformational annealing upon rf heating on a second timescale applicable in TIMS and FAIMS, but not DTIMS. While TIMS could be operated in a “soft” regime with minimal heating and conformational distortion of proteins,^43^ that (unlike in native MS studies) is irrelevant here.

In a set of five variants making 10 pairs, we separated four pairs fully and three in part. The results were confirmed for precursors employing mixtures and at the fragment level by unique PTM-containing products of collision-induced dissociation. This performance is more useful than appears as “partial separations” filtering one variant allow detecting and quantifying the other by balance.^44^ Hence, actually 70% of variant pairs considered here could be fully disentangled. Also, as unresolved binary mixtures can be characterized by unique fragments, only separation into those rather than single species is truly necessary.

Better variant resolution is still desired. As long as the individual conformers are in effect separated as above, that could be reached by raising the instrumental resolving power. Novel cyclic IMS (cIM) platforms that involve rf heating of similar magnitude and duration but reach R_IMS_ > 500 may deliver the solution.^30^ An alternative or complementary avenue is 2-D IMS, in the FAIMS/IMS configuration^45^ or tandem linear IMS^46,47^ with ions activated between the stages to create orthogonality. While ECD and ETD are suitable with all PTMs and provide broader sequence coverage, they were hampered by low yield, long reaction times incompatible with short transients from dispersive IMS methods such as TIMS or cIMS, and the FTICR requirement for ECD. With those obstacles now largely overcome by rapid “hot ECD” in electromagnetostatic cells,^48,49^ augmenting the present workflow by ECD is straightforward.

The now enabled capability for intact proteoform separations is most promising for top-down analyses of modified proteomes in general and in the epigenetic context in particular.

## Supporting information

Supplemental results

## ACKNOWLEDGMENTS

This study was supported by grants to O.N.J. from the Independent Research Fund Denmark (0135-00114B) and the Novo Nordisk Foundation (INTEGRA, NNF20OC0061575), and the NSF CAREER award to A.S. (CHE-1552640). We thank Neil Kelleher (Northwestern) and Benjamin Garcia (Washington U.) for insightful discussions.

## Supporting Information Available

Detailed description of histone synthesis. Table showing inverse mobilities, collision cross section and resolving power values for H4K5ac, H4K8ac, H4K12ac, H4K16ac and H4K20ac. Additional TIMS spectra for isomeric pairs of H4ac proteoforms and charge states not presented in Figure 3 for z = 16 – 19.

