## Supplemental results for "Top-Down Ion Mobility Separations of Isomeric Proteoforms"

**Scheme S1 - Reaction scheme and methods – Pages S2**

**Histone Synthesis method – Pages S3-4**

**Figure S1 – Pages S5**

**Figure S2 – Pages S6**

**Figure S3– Pages S7**

**Table S1 – Page S8**

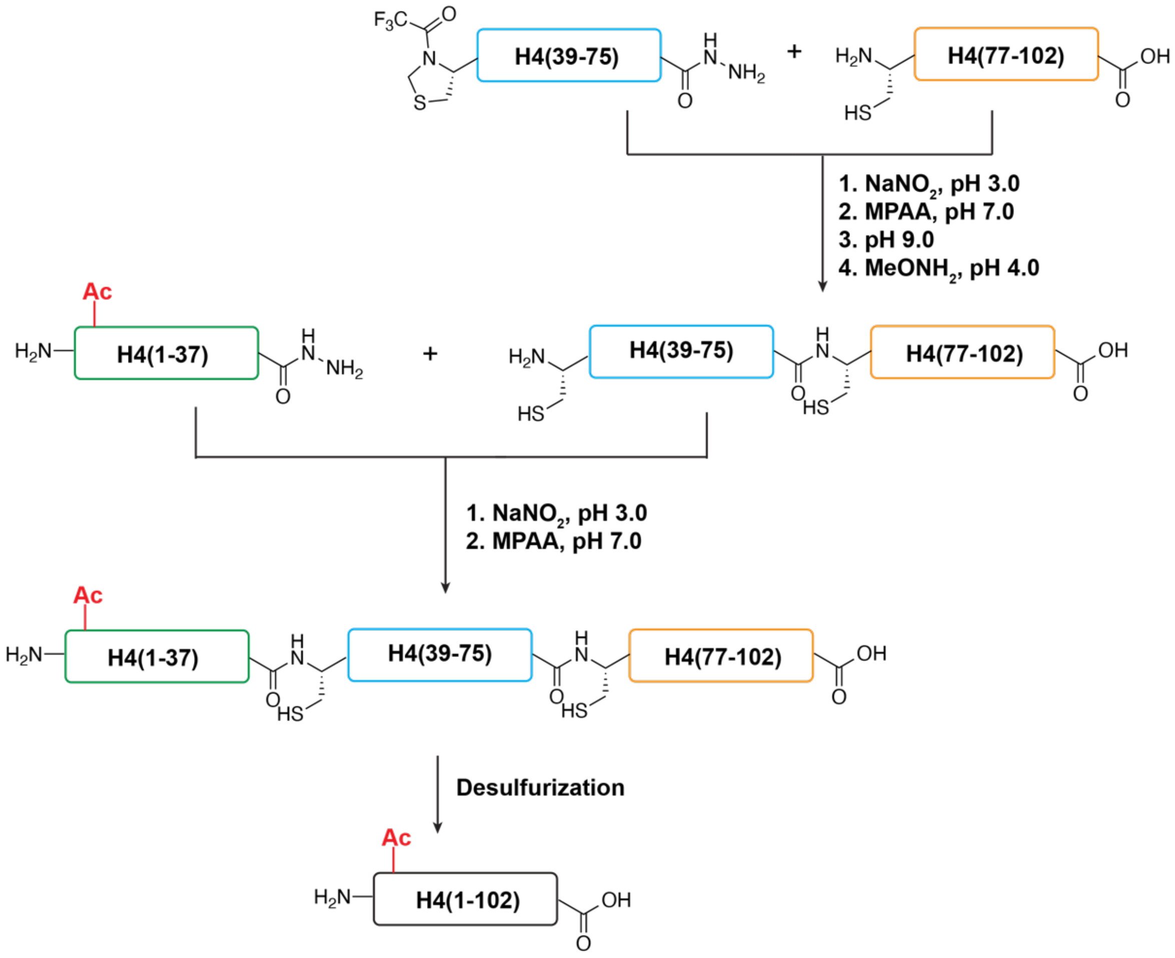


**Scheme S1**. Synthesis of full-length H4 proteins with acetyllysine.

### Histone synthesis method

#### General information

All Fmoc protected amino acids, HCTU, DIEA, MPAA, TFA, 4-methylpiperidine, TIPS, and Fmoc-Gly-Wang resin were purchased from Chemimpex Inc. 2-chlorotrityl resin was purchased from Chempep Inc. VA-044 and TECP were purchased from TCI Chemicals. HPLC grade acetonitrile was purchased from VWR. All other reagents were purchased from Fisher Scientific.

#### Preparation of hydrazine resin.

The hydrazine resin was freshly prepared before peptide synthesis following a previously published procedure4. Briefly, 200 mg 2-chlorotrityl resin was swelled in 2 mL DMF for 15 min. A mixture of DIEA (125 mL) and 10% hydrazine hydrate in DMF (380 mL) was added dropwise to the resin, and the suspension was shaken at room temperature for 2 h. 2 mL MeOH was then added to the suspension, which was shaken for additional 15 min. Finally, the resin was washed with DMF (3 mL, x4).

The C-terminal amino acid was introduced to the hydrazine resin by treating resin with Fmoc-AA (4 eq.), DIEA (16 eq.) and HCTU (3.8 eq) in 2 mL DMF for 1 h at room temperature. The coupling reaction was repeated.

#### Fmoc-based solid-phase peptide synthesis.

*Synthesis of peptidyl hydrazide.* Solid-phase peptide synthesis (SPPS) was conducted on a Purepep Chorus peptide synthesizer with the pre-loaded hydrazine resin (0.1 mmol scale). A typical SPPS cycle includes Fmoc deprotection (3 mL 20% 4-methylpiperidine/DMF, 10-min shaking at rt, repeated), DMF wash (4 mL, x3), coupling (4 eq. Fmoc-AA, 3.8 eq. HCTU, 8 eq. DIEA in 4 mL DMF, 120-min shaking at RT), and another DMF washing step (4 mL, x3). After SPPS cycles were completed, a final Fmoc deprotection step was conducted.

*Synthesis of the C-terminal segment.* Solid-phase peptide synthesis (SPPS) was conducted on a Purepep Chorus peptide synthesizer with the pre-loaded wang resin (0.1 mmol scale). A typical SPPS cycle includes Fmoc deprotection (3 mL 20% 4-methylpiperidine/DMF, 10-min shaking at rt, repeated), DMF wash (4 mL, x3), coupling (4 eq. Fmoc-AA, 3.8 eq. HCTU, 8 eq. DIEA in 4 mL DMF, 10-min shaking at 75 °C), and another DMF washing step (4 mL, x3). After SPPS cycles were completed, a final Fmoc deprotection step was conducted.

*Peptide cleavage.* The resin was washed with DMF (4 mL, x3) and DCM (4 mL, x5) and dried with N2 flow for 20 min. Peptide cleavage was done using 5 mL TFA cleavage cocktail (TFA:TIPS:H2O:DTT, 94:2:2:2) for 3 h. The resin slurry was filtered, and the filtrate was added dropwise to 45 mL cold diethyl ether (pre-chilled on dry ice). A white precipitate was collected by centrifugation and washed with 40 mL cold ether. The crude peptide was obtained after air drying.

#### General procedure of HPLC purification

The crude peptide was dissolved in 10 mL 6 M GnHCl and filtered through a 0.45 mm syringe filter. The crude material was purified by reverse-phase HPLC using a C4 preparative column (Higgins Analytical, Inc.) and solvent A (water, 0.1% TFA)/B (ACN, 0.1% TFA) as the mobile phase.

#### Hydrazide-based native chemical ligation

*Synthesis of Cys-H4(39-75)-Cys-H4(77-102)-OH.* Tfa-Thz-H4(39-75)-NHNH2 (10 mg) was dissolved in an activation buffer (6 M GnHCl, 100 mM sodium phosphate, pH 3.0) to 2 mM. The solution was sonicated for 10 min and chilled in a salt/ice bath for 10 min. The activation of hydrazide was done by adding freshly prepared $NaNO_2$ solution (0.3 M in the activation buffer) to reach a final concentration of 30 mM. The mixture was incubated in a salt/ice bath for 20 min. Then, an equal volume of a freshly prepared MPAA solution (0.2 M MPAA in 6 M GnHCl, 100 mM sodium phosphate, pH 7.5) was added to the mixture. The resulting solution was allowed to warm up to room temperature and added to Cys-H4(77-102)-OH (7 mg). After 10-min sonication, the final pH was adjusted to 6.8-7.0 by the addition of 2 M NaOH (increment of 2 mL). The solution was kept at rt for overnight. Upon completion, the pH was adjusted to 9.0 with 2 M NaOH. After 1-hour incubation at room temperature, $MeONH_2$·HCl was added to a final concentration of 0.3 M, and the solution pH was adjusted to 4.0 with 2 M HCl. After reaction for 4 h at room temperature, an equal volume of TCEP solution (0.1 M in 6 M GnHCl, pH 7) was added to the reaction mixture, and the solution was incubated for 10 min. The solution was acidified by mixing with TCEP solution (0.1 M in 6 M GnHCl, pH 2). After centrifugation at 14,000 g for 10 min, the supernatant was collected, purified by reverse-phase HPLC, and freeze-dried to give Cys-H4(39-75)-Cys-H4(77-102)-OH.

*Synthesis of H4(1-37)-Cys-H4(39-75)-Cys-H4(77-102)-OH*. H4(1-37)-$NHNH_2$ (1 mg) was dissolved in an activation buffer (6 M GnHCl, 100 mM sodium phosphate, pH 3.0) to 2 mM. The solution was sonicated for 10 min and chilled in a salt/ice bath for 10 min. The activation of hydrazide was done by adding freshly prepared $NaNO_2$ solution (0.3 M in the activation buffer) to reach a final concentration of 30 mM. The mixture was incubated in a salt/ice bath for 20 min. Then, an equal volume of a freshly prepared MPAA solution (0.2 M MPAA in 6 M GnHCl, 100 mM sodium phosphate, pH 7.5) was added to the mixture. The resulting solution was allowed to warm up to room temperature and added to Cys-H4(39-75)-Cys-H4(77-102)-OH (2 mg). After 10-min sonication, the final pH was adjusted to 6.8-7.0 by the addition of 2 M NaOH (increment of 2 mL). After reaction for overnight at room temperature, an equal volume of TCEP solution (0.1 M in 6 M GnHCl, pH 7) was added to the reaction mixture, and the solution was incubated for 10 min. The solution was acidified by mixing with TCEP solution (0.1 M in 6 M GnHCl, pH 2). After centrifugation at 14,000 g for 10 min, the supernatant was collected, purified by reverse-phase HPLC, and freeze-dried to give H4(1-37)-Cys-H4(39-75)-Cys-H4(77-102)-OH.

Desulfurization. H4(1-37)-Cys-H4(39-75)-Cys-H4(77-102)-OH (2 mg) was dissolved in 1 mL desulfurization buffer (0.2 M TCEP, 0.2 M NaPhos, 8 M GnHCl, pH 7.0, degassed under vacuum). VA-044 (7 mg) and reduced glutathione (12.5 mg) were added, and the pH was adjusted to 6.8 with 4 M NaOH. The resulting mixture was incubated at 37 °C for overnight. After the completion of desulfurization, acetic acid was added to 5% (v/v). After centrifugation at 14,000 g for 10 min, the supernatant was collected, purified by reverse-phase HPLC, and freeze-dried to give full-length H4 proteins.

1. Li, J.; Li, Y.; He, Q.; Li, Y.; Li, H.; Liu, L., One-pot native chemical ligation of peptide hydrazides enables total synthesis of modified histones. *Org Biomol Chem* **2014,** *12*, 5435-41.
2. Huang, Y. C.; Chen, C. C.; Gao, S.; Wang, Y. H.; Xiao, H.; Wang, F.; Tian, C. L.; Li, Y. M., Synthesis of l- and d-Ubiquitin by One-Pot Ligation and Metal-Free Desulfurization. *Chemistry* **2016,** *22* (22), 7623-8.
3. Fang, G. M.; Li, Y. M.; Shen, F.; Huang, Y. C.; Li, J. B.; Lin, Y.; Cui, H. K.; Liu, L., Protein chemical synthesis by ligation of peptide hydrazides. *Angew Chem Int Ed Engl* **2011,** *50*, 7645-9.
4. Zheng, J. S.; Tang, S.; Qi, Y. K.; Wang, Z. P.; Liu, L., Chemical synthesis of proteins using peptide hydrazides as thioester surrogates. *Nat Protoc* **2013,** *8*, 2483-95.

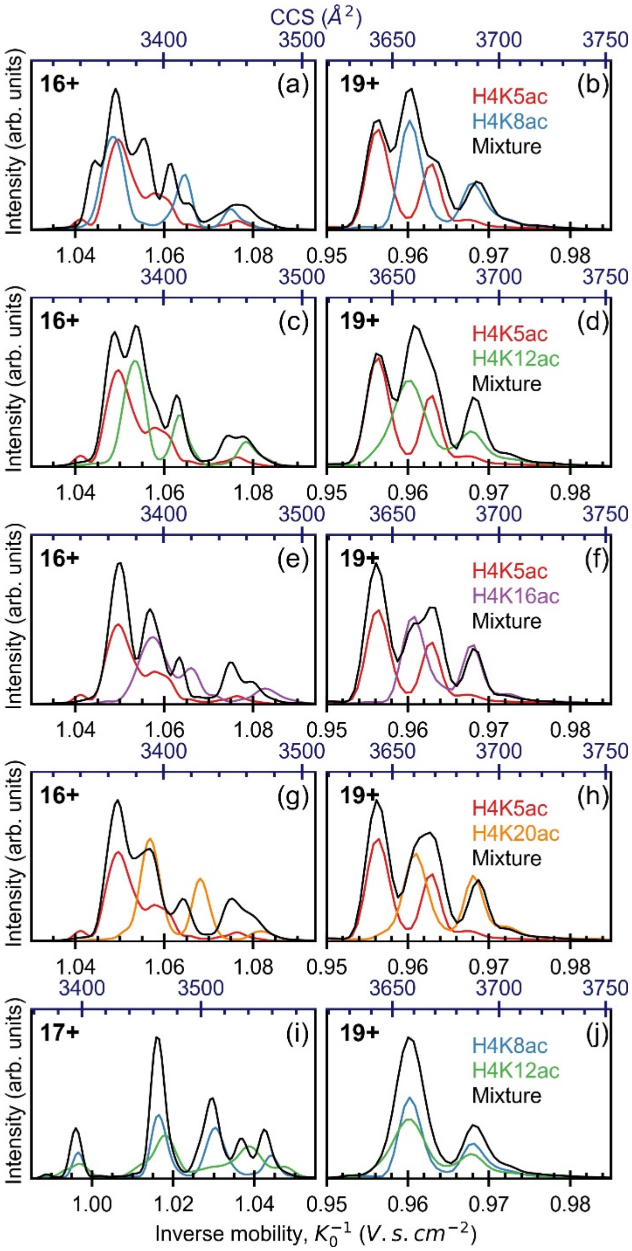


**Figure S1.** TIMS-MS separation of binary mixture (black) of H4K5ac (red), H4K8ac (blue), H4K12ac (green), H4K16ac (violet) and H4K20ac (orange) for charge state 16, 17 and 19+. Experiments were performed using $P_{in}$ = 3.4 mbar and an electric field scan rate of 17.5 V/s.

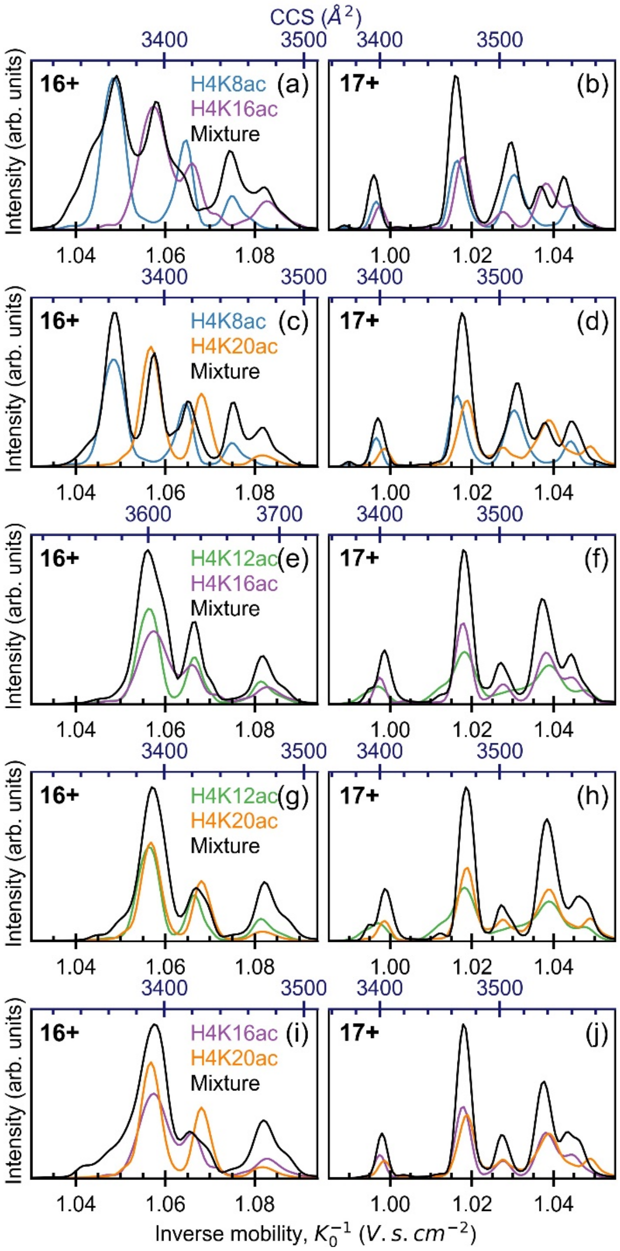


**Figure S2.** TIMS-MS separation of binary mixture (black) of H4K8ac (blue), H4K12ac (green), H4K16ac (violet) and H4K20ac (orange) for charge state 16 and 17+. Experiments were performed using $P_{in}$ = 3.4 mbar and an electric field scan rate of 17.5 V/s.

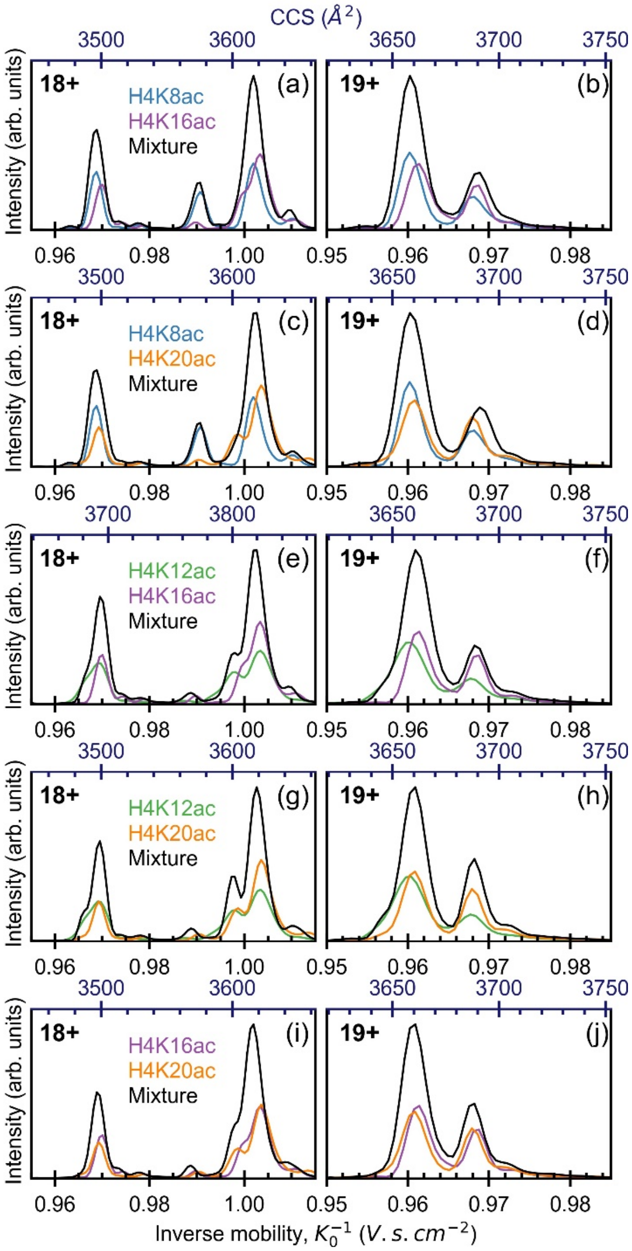


**Figure S3.** TIMS-MS separation of binary mixture (black) of H4K8ac (blue), H4K12ac (green), H4K16ac (violet) and H4K20ac (orange) for charge state 18 and 19+. Experiments were performed using $P_{in}$ = 3.4 mbar and an electric field scan rate of 17.5 V/s.

**Table S1. Average inverse mobility (<K0$^{-1}$>), average collision cross section (CCS) and resolving power (R) of positional PTM isomers of acetylated H4 (H4KXac, with X = 5, 8, 12, 16 and 20)**

| z | Histone | <K0-1> | CCS ($A^2$) | R |
|---|---|---|---|---|
| 19 | H4K5ac | 0.9563 | 3643 | 288 |
| | | 0.9629 | 3668 | 320 |
| | H4K8ac | 0.9603 | 3658 | 194 |
| | | 0.9679 | 3687 | 213 |
| | H4K12ac | 0.9601 | 3657 | 194 |
| | | 0.9679 | 3687 | 213 |
| | H4K16ac | 0.9614 | 3662 | 270 |
| | | 0.9685 | 3690 | 273 |
| | H4K20ac | 0.9573 | 3647 | 273 |
| | | 0.9642 | 3673 | 303 |
| 18 | H4K5ac | 0.9642 | 3480 | 323 |
| | | 0.9965 | 3597 | 324 |
| | H4K8ac | 0.9687 | 3496 | 317 |
| | | 0.9907 | 3576 | 294 |
| | | 1.0020 | 3616 | 260 |
| | | 1.0101 | 3646 | 243 |
| | H4K12ac | 0.9688 | 3497 | 172 |
| | | 0.9972 | 3599 | 183 |
| | | 1.0036 | 3622 | 177 |
| | H4K16ac | 0.9699 | 3500 | 311 |
| | | 1.0029 | 3620 | 232 |
| | | 1.0111 | 3649 | 163 |
| | H4K20ac | 0.9693 | 3498 | 295 |
| | | 0.9980 | 3602 | 309 |
| | | 1.0038 | 3623 | 193 |
| 17 | H4K5ac | 0.9915 | 3379 | 323 |
| | | 1.0109 | 3446 | 242 |
| | | 1.0209 | 3480 | 231 |
| | | 1.0328 | 3520 | 189 |
| | H4K8ac | 0.9966 | 3397 | 322 |
| | | 1.0166 | 3465 | 219 |
| | | 1.0305 | 3512 | 176 |
| | | 1.0441 | 3559 | 217 |
| | H4K12ac | 0.9963 | 3396 | 149 |
| | | 1.0180 | 3470 | 163 |
| | H4K16ac | 0.9975 | 3400 | 318 |
| | | 1.0178 | 3469 | 233 |
| | | 1.0272 | 3501 | 247 |
| | | 1.0383 | 3539 | 169 |
| | | 1.0459 | 3565 | 251 |
| | H4K20ac | 0.9986 | 3404 | 270 |
| | | 1.0187 | 3472 | 212 |
| | | 1.0385 | 3540 | 151 |
| | | 1.0490 | 3575 | 197 |
| 16 | H4K5ac | 1.0496 | 3367 | 168 |
| | | 1.0583 | 3395 | 129 |
| | H4K8ac | 1.0484 | 3363 | 178 |
| | | 1.0643 | 3414 | 232 |
| | | 1.0749 | 3448 | 236 |
| | H4K12ac | 1.0819 | 3471 | 169 |
| | | 1.0562 | 3388 | 190 |
| | | 1.0814 | 3469 | 223 |
| | H4K16ac | 1.0662 | 3420 | 182 |
| | | 1.0573 | 3392 | 142 |
| | | 1.0828 | 3474 | 158 |
| | H4K20ac | 1.0568 | 3390 | 197 |
| | | 1.0682 | 3427 | 229 |
| | | 1.0820 | 3471 | 149 |
